# Similarity-weighted entropy for quantifying genetic diversity in viral quasispecies

**DOI:** 10.1101/2024.10.06.616857

**Authors:** Jian Wu

## Abstract

Viral quasispecies are dynamic populations of genetically diverse viruses, often exhibiting high mutation rates. Understanding the genetic diversity within these quasispecies is critical for analyzing viral evolution, adaptation, and treatment resistance. Entropy and normalized Shannon entropy are widely used metrics to quantify this diversity. However, these metrics ignore genetic similarities between sequences, potentially underestimating the true diversity. In this paper, we introduce two methods for similarity-weighted normalized entropy that account for sequence similarities and provide more accurate measures of genetic diversity. By applying these methods to two hypothetical viral quasispecies populations, we compare the traditional entropy, normalized entropy, and the proposed similarity-weighted measures. Our results demonstrate that the similarity-weighted entropies better capture the true genetic diversity in highly related viral populations, while retaining the simplicity of the original entropy calculations. We discuss the advantages and limitations of both similarity-weighted measures and propose their application in viral quasispecies studies.

## 1. Introduction

Viral quasispecies, originally proposed by Eigen in 1971 (Eigen 1971), represent a population of related viral sequences within an infected host (Domingo and Perales 2019). Due to the error-prone nature of viral replication, especially in RNA viruses and viroids, a high degree of genetic diversity exists among these sequences (Gago et al. 2009; Jones et al. 2021; Wu and Bisaro 2020). This diversity allows viral quasispecies to adapt rapidly to changing environments, evade immune responses, and develop resistance to antiviral treatments (Domingo et al. 2021).

Quantifying genetic diversity is essential for understanding the behavior and evolution of quasispecies. Shannon entropy and normalized entropy are commonly used to describe the genetic heterogeneity of a population, with higher entropy reflecting greater diversity (Arbiza et al. 2010; Gregori et al. 2016; Gregori et al. 2023; Wu et al. 2024). However, traditional entropy measures do not take into account the genetic similarity between sequences. In populations where sequences differ by only a few mutations, entropy may overestimate the effective diversity by treating similar sequences as entirely distinct. This limitation calls for an improved metric that incorporates sequence similarity into entropy calculations. Weighted entropy is a variation of the standard Shannon entropy, where different elements in a dataset contribute differently based on assigned weights (Guiaşu 1971; Kelbert et al. 2017; Mahdy 2018). In a biological or genetic context, weighted entropy can be used to measure the diversity or uncertainty of sequences while accounting for the importance or similarity of individual sequences (Chang and Wang 2011; Xie et al. 2024). However, the use of weighted entropy to analyze the genetic diversity of viral quasispecies based on sequence similarity has not yet been reported.

In this study, we propose two methods for calculating similarity-weighted normalized entropy. These new measures aim to correct for the neglect of sequence similarity in traditional entropy metrics. We demonstrate their application in two hypothetical viral quasispecies populations and compare the results to standard entropy measures. Our analysis highlights the advantages of incorporating similarity to better reflect the true genetic diversity of viral quasispecies.

## 2. Methods

### 2.1 Traditional Shannon entropy and normalized Shannon entropy

Shannon entropy *H* is calculated using:

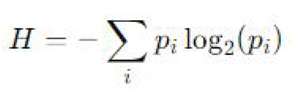

where *pi* is the frequency of the *i*-th sequence in the population. Normalized entropy *H*n is derived as:

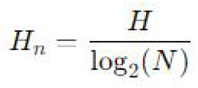

### 2.2 Similarity-weighted normalized Shannon entropy (Method 1)

The first new measure introduces a similarity-weighted factor to the normalized Shannon entropy:

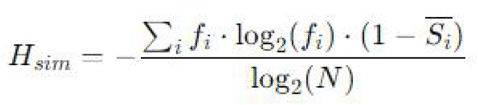

Here, *fi* is the frequency of sequence *i*, and 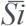 is the average similarity of sequence *i* to all other sequences.

### 2.3 Pairwise similarity-weighted normalized Shannon entropy (Method 2)

The second approach incorporates all pairwise sequence similarities:

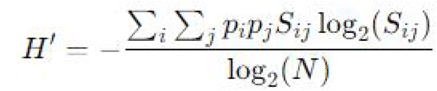

where *S*_*ij*_ represents the similarity between sequences *i* and *j*, and *pi* and *pj* are their respective frequencies.

### 2.4 Sequence data and similarity matrices

We constructed two hypothetical viral quasispecies populations, each with three sequences and the same frequency distribution, but different similarity matrices:

Population A Frequencies: 40% (A), 30% (B), 30% (C)

Population B Frequencies: 40% (A), 30% (B), 30% (C)

Population A similarity matrix:

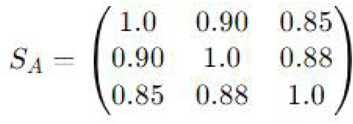

Population B similarity matrix:

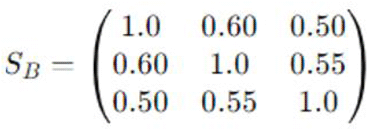

## 3. Results

### 3.1. Traditional entropy and normalized entropy

Both populations have the same sequence frequencies, yielding identical values for traditional Shannon entropy and normalized Shannon entropy.

Shannon entropy for both population A and population B:

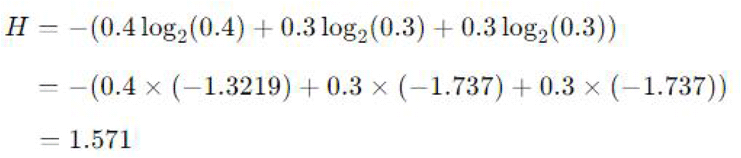

Normalized Shannon entropy for both populations:

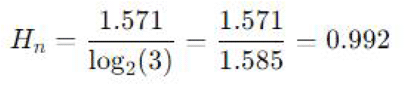

### 3.2 Similarity-weighted normalized Shannon entropy (Method 1)

Step 1: Calculate the average similarity (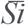 )for each sequence in population A population B.

Population A:

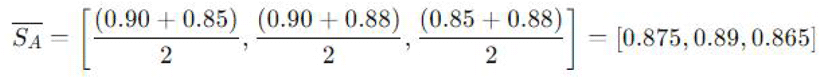

Similarly, population B:

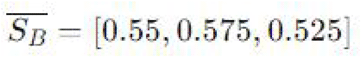

Step 2: Apply the similarity-weighted Shannon entropy equation for population A and population B.

Population A:

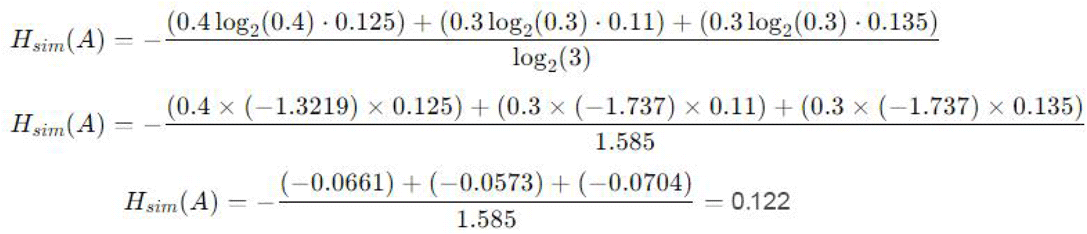

Population B:

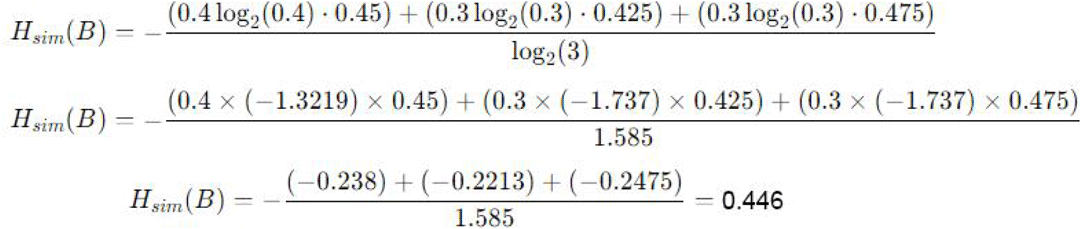

### 3.3 Pairwise similarity-weighted normalized Shannon entropy (Method 2)

For population A:

Step 1: Calculate weighted contributions for each Pair

For each pair of sequences, we calculate *pipjS*_*ij*_log_2_(*S*_*ij*_). Note that for *i*=*j, Sij*=1, so log_2_(*S*_*ij*_)=0, and the contribution is zero.

For *S*_*11*_ : *S*_*11*_=1, so log_2_(1)=0, and 0.4×0.4×1×log_2_(1)=0

For *S*_*12*_ : *S*_*12*_=0.9, so log_2_(0.7)≈−0.152, and 0.4×0.3×0.9×log_2_(0.7)=−0.0164

Similarly, the values for *S*_*13*_, *S*_*21*,_ *S*_*22*_, *S*_*23*_, *S*_*31*_, *S*_*32*_ and *S*_*33*_ are −0.0239, −0.0164, 0, −0.0152, -0.0239, −0.0152, and 0, respectively. Step 2: Sum of all pairwise terms

The sum of the non-zero contributions:

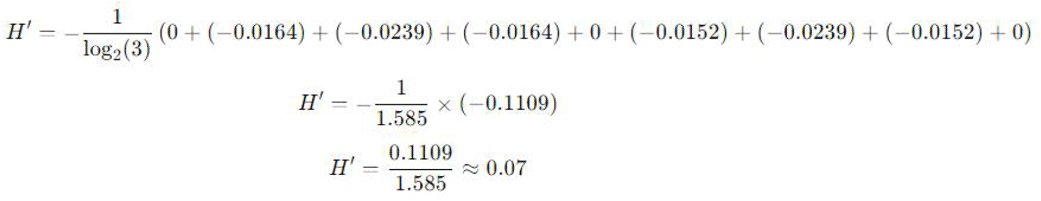

For population B:

Similarly, the values for *S*_*11*_, *S*_*12*_, *S*_*13*_, *S*_*21*,_ *S*_*22*_, *S*_*23*_, *S*_*31*_, *S*_*32*_ and *S*_*33*_ are 0, -0.0531, -0.06, -0.0531, 0, −0.0425, −0.06, −0.0425 and 0, respectively.

Step 2: Sum of all pairwise terms

The sum of the non-zero contributions:

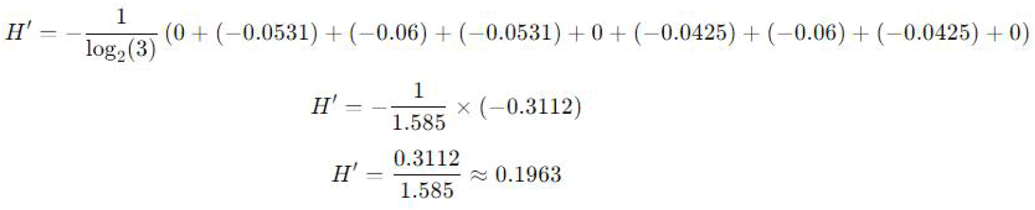

### 3.4 Summary of results

Traditional Shannon entropy and normalized Shannon entropy (for both populations):

*H*=1.571, *H*_*n*_=0.992

Similarity-weighted Shannon entropy (Method 1): Population A:

*H*_*sim*_(*A*)=0.122

Population B:

*H*_*sim*_(*B*)=0.446

Pairwise similarity-weighted normalized Shannon entropy (Method 2)

Population A:

*H*’(*A*)=0.07

Population B:

*H*’(*B*)=0.1963

## 4. Discussion

The results demonstrate significant differences between the two approaches to similarity-weighted entropy. Although both populations have identical traditional entropy values, the incorporation of sequence similarity reveals the underlying genetic structure more accurately.

Traditional Shannon entropy provides a general measure of genetic diversity by considering sequence frequencies alone (Gregori et al. 2014; Smerlak 2021). In both quasispecies populations, the normalized entropy (*H*=0.992) suggests a high degree of diversity. However, this metric neglects the fact that many sequences may be closely related, particularly in viral quasispecies where mutations are often incremental (Brass et al. 2017; Miller et al. 2011; Smith et al. 1997; Wu et al. 2020; Wu and Bisaro 2024).

The first method, which weights entropy by the average similarity of each sequence, demonstrates a more nuanced view of diversity. For Population A, which exhibits high similarity between sequences (average similarity values around 85-90%), the diversity is lower than suggested by traditional entropy, yielding *H*_*sim*_ (A)=0.122. In contrast, Population B, with lower average similarities (around 50-60%), shows a higher similarity-weighted entropy *H*_*sim*_(*B*)=0.446, indicating that the sequences are more genetically distinct from one another. This method accounts for both the frequency and the structural relatedness of sequences, providing a more refined assessment of genetic diversity.

The second approach, based on pairwise sequence similarity, produces even lower entropy values for Population A (*H’*(*A*)=0.07), which reflects the high similarity across all pairs of sequences. For Population B, the pairwise method yields *H’*(*B*)=0.1963, again showing more diversity than in Population A, but less than the result from Method 1. This method directly integrates the similarity between each pair of sequences, making it more sensitive to small differences across the population. This makes it a powerful tool for comparing populations with subtle differences in their genetic makeup.

Each method offers unique insights into the genetic diversity of viral quasispecies. Method 1 (average similarity weighting) smooths the diversity calculation by considering the overall similarity of each sequence to the population. It is computationally simpler and may be more intuitive when the goal is to capture the general similarity structure. This method balances computational simplicity with a reasonable reflection of genetic diversity, making it suitable for large viral populations or initial diversity assessments. However, it can sometimes overlook subtle pairwise relationships, potentially underestimating diversity when some sequence pairs are highly divergent. Method 2 (pairwise similarity weighting) is more granular, accounting for the relationships between all sequence pairs. This method provides a more detailed picture of the population’s genetic structure, but may be more computationally intensive, particularly for large datasets. This method offers higher precision by directly comparing all sequences. This makes it a better choice for small to medium-sized populations or when exact relationships between sequences are of primary interest. The downside is that it requires more computational resources and may become inefficient for very large quasispecies populations.

## 5. Conclusion

This study introduced two new methods to calculate similarity-weighted normalized entropy for analyzing viral quasispecies diversity. By incorporating sequence similarity, these methods provide a more accurate picture of the genetic structure of viral populations, particularly when traditional entropy measures may overestimate diversity. Both approaches offer valuable insights, with the choice of method depending on the specific requirements of the analysis. These methods can potentially improve the study of viral evolution and the development of strategies for managing viral infections.

## Data availability

All the data are presented in this paper.

### Funding

This study was supported by the National Natural Science Foundation of China (32272483, 32470149).

